# Phage-encoded CasPRs transcriptionally silence diverse CRISPR-Cas systems

**DOI:** 10.64898/2026.02.23.707548

**Authors:** Edith M. Sanderson, Julieta Peralta, Sam Nouwens, Luke Oriolt, Victoria M. Hayes, Brett K. Kaiser, Alexander J. Meeske

## Abstract

Anti-CRISPRs (Acrs) are diverse proteins or RNAs that protect invading phages and plasmids from host CRISPR-Cas immunity. Most Acrs neutralize their cognate Cas proteins via direct physical interaction. Here we describe CasPRs, a particularly widespread family of DNA-binding Acrs that recognize specific sequence motifs within cas gene coding regions, thereby blocking RNA polymerase and silencing transcription. We demonstrate that eight diverse CasPRs bind to the *cas8b* gene to repress the type I-B CRISPR-Cas system in its native host, *Listeria seeligeri*. Meanwhile, a CasPR from *Streptococcus dysgalactiae* silences type II-A CRISPR-Cas immunity by binding to the *cas9* coding sequence. We found that one CasPR is required to inhibit CRISPR immunity during lysogeny by its host prophage. Taken together, our results indicate that members of the CasPR family have diverged to silence completely unrelated CRISPR types, and suggest transcriptional repression is a common mode of phage-mediated immune antagonism.

## Introduction

CRISPR-Cas systems protect their prokaryotic hosts from viral infection using RNA-guided Cas nucleases that target and cleave foreign nucleic acids in a sequence-specific manner^1-7^. Bioinformatic analyses over the past 20 years have revealed millions of diverse CRISPR loci, which are classified into seven types that differ in their nuclease content and mechanism of target interference^8^. To subvert CRISPR immunity during infection, phage encode anti-CRISPRs (Acrs), which are proteins or RNAs that inhibit Cas nuclease function^9,10^. Typical Acrs interact with specific Cas proteins, have a limited inhibitory range that does not span across multiple CRISPR types, and are named after the sub-type of CRISPR they antagonize^11,12^.

Acrs inhibit CRISPR immunity via diverse mechanisms, but usually work post-translationally to neutralize the activities of Cas nucleases. Many Acrs bind cognate Cas proteins and block their interaction with target nucleic acids^13^. In contrast, a small subset of Acrs have been shown to affect the steady-state levels of Cas interference complexes. AcrIIA1, which inhibits the single effector nuclease Cas9, forms a protein-protein interaction with Cas9 and causes degradation of the protein via an unknown mechanism^14^. AcrIF25 disassembles the fully formed multi-subunit type I-F Cascade interference complex^15^. AcrVA2 binds to the N-terminus of the nascent Cas12a polypeptide, causing degradation of the *cas12a* mRNA in response, and to date is the sole Acr protein demonstrated to affect Cas protein biogenesis^16^.

Genes encoding Acrs are often clustered together within the genomes of phages, prophages, and plasmids. These “inhibitor islands” often contain genes with diverse inhibition specificity corresponding to the repertoire of immune systems the invading element is likely to encounter. They commonly harbor multiple acr gene families and inhibitors of restriction-modification systems^17^. In addition, many such clusters contain helix-turn-helix (HTH) domain-containing transcription factors called anti-CRISPR-associated proteins (Acas), which perform autorepression of acr gene expression that is otherwise detrimental to the phage transcriptional program during infection^18^.

Here we report a widespread group of phage-encoded DNA-binding proteins that act as Acrs; they recognize specific sites in cas gene coding sequences and silence transcription to prevent biogenesis of CRISPR-Cas machinery. Genes encoding these factors are defined by the presence of both CapR and AP2-family DNA-binding domains and are highly abundant in acr loci within phages of phylum Bacillota. Working in *Listeria seeligeri*, a native host of both type I-B and II-A CRISPR systems, we characterized eight representatives of this family that silence the type I-B (Cascade) system and one that silences the type II-A (Cas9) system. As these are members of a single protein family that have diverged in evolution to inhibit completely unrelated CRISPR types, they defy the established nomenclature for Acrs. Accordingly, we refer to them as CasPRs (Cas Protein Repressors).

## Results

### Two Acr proteins affect type I-B cas gene expression

We previously screened a collection of 62 *Listeria seeligeri* strains for anti-CRISPR activities against four *Listeria* CRISPR types (I-B, II-A, II-C, VI-A) arising from endogenous prophages and plasmids residing within each strain^11^. By testing individual genes located within putative acr operons in each strain for the ability to inhibit the relevant CRISPR type, we identified the founding members of 12 Acr protein families. Of these, the largest class (8 out of 12, AcrIB3-10) inhibits the *L. seeligeri* str. LS1 type I-B CRISPR-Cas system (**Fig. 1A**), which is predominant among *Listeria spp*. While this system is transcriptionally silent under laboratory growth conditions, it potently restricts targeted plasmids when placed under the control of a constitutive Ptet promoter on a single-copy integrating plasmid^11,19^. We continued this Acr discovery effort, uncovering six additional unique proteins that relieve plasmid targeting by the type I-B system, here called AcrIB11-16, for a total of 14 type I-B inhibitor families (**Fig. 1B-C and S1A-B**). While we previously investigated the mechanisms of AcrIB3 and IB4, the other 12 remain uncharacterized^11^. We began by testing whether these Acrs affect the levels of the type I-B nuclease Cas3. We introduced plasmids with each Acr into strain LS1 harboring the type I-B system with a functionally tagged Cas3-3xFlag, and immunoblotted for Cas3 (**Fig. 1D**). Of the twelve Acrs tested, ten had no effect on Cas3 protein levels, however in the presence of AcrIB12 or AcrIB14, Cas3 was undetectable. Thus, unlike most Acrs, these two proteins affect the biogenesis or stability of CRISPR-Cas machinery, and the remainder of our analysis focuses on them exclusively.

**Figure 1.**
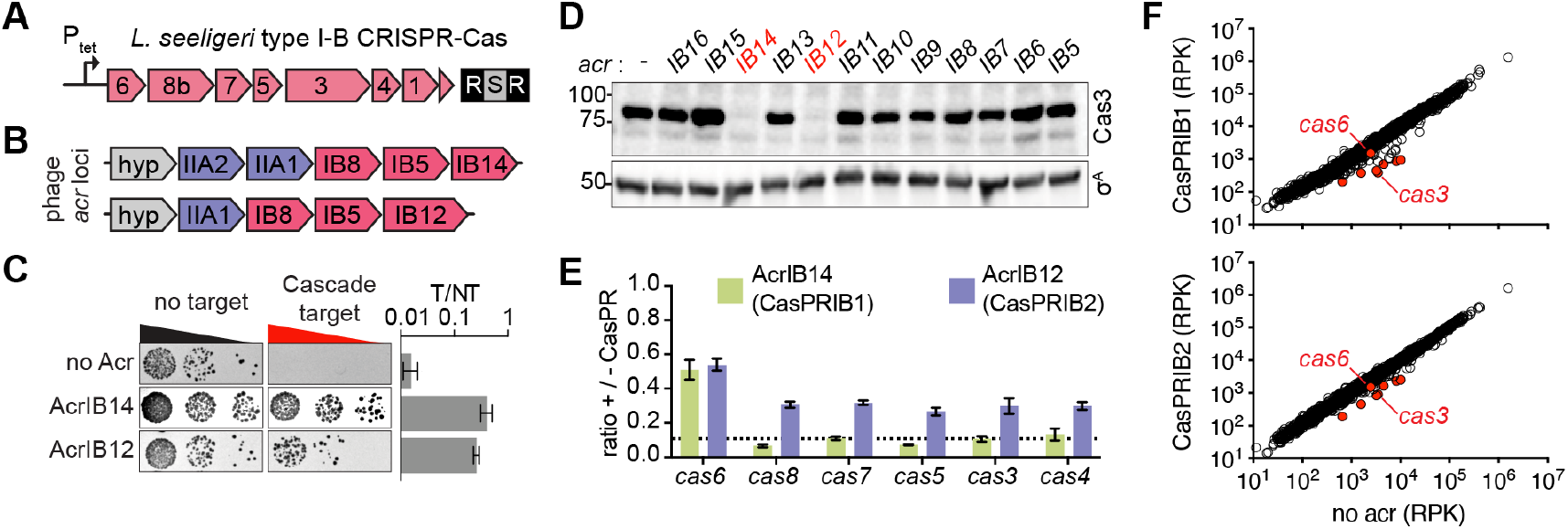
CasPRs affect expression of type l-B cas genes. **(A)** Schematic showing chromosomally integrated *L. seeligeri* str. LS1 type l-B CRISPR-Cas locus with single plasmid-targeting spacer under Ptet control. **(B)** Prophage acr loci from *L. seeligeri* str. RR4 (top) and str. FSL W9-0499 (bottom) showing type Il-A (blue) and type l-B (pink) acr genes. **(C)** Plasmid targeting assay showing AcrlB14 and AcrlB12 inhibition of type l-B CRISPR interference. T/NT, ratio of target to non-target transconjugants. Error bars represent SEM. **(D)** Anti-Flag western blot detecting Cas3-3xFlag and σ ^A^ loading control in the presence of the indicated type l-B Acrs. Molecular weight standards shown in kDa. Acts highlighted in red cause diminished Cas3 levels. **(E)** RT-qPCR experiments measuring cas gene transcripts during Acr expression. Expression values shown are relative to a no-Acr control, set to 1. Ct values were normalized to expression of the housekeeping gene ssbA. Dashed horizontal line indicates limit of detection. Error bars represent SEM. (F) RNA-seq comparing reads per kilobase (RPK) for each *L. seeligeri* gene in presence of CasPRIBI (top, y-axis), CasPRIB2 (bottom, y-axis), or empty vector control (x-axis). Type l-B cas genes highlighted in red. Each point represents mean of 3 biological replicates.

AcrIB12 and IB14 are each found in *L. seeligeri* prophage *acr* loci alongside other type I-B and type II-A *acr* gene homologs (**Fig. 1B**). Upon closer inspection of the AcrIB12 and IB14 amino acid sequences, we determined that they are distantly related, as 40% of the AcrIB12 protein shares 55% identity with AcrIB14, suggesting they may share an inhibition mechanism (**Fig. S1C**). Next, we performed RT-qPCR on each cas gene along the type I-B locus to investigate whether AcrIB12 or IB14 affect the levels of *cas* gene transcripts across the entire operon (**Fig. 1E**). During Acr expression, we detected between 2- and 10-fold depletion for all *cas* gene transcripts, often falling below the limit of detection. The depletion effect was stronger for AcrIB14 than for AcrIB12 and was less potent on *cas6* (the first gene in the operon) than for all other *cas* genes. We note that since transcription of the type I-B system is driven by a synthetic promoter in this experiment, these factors are unlikely to repress expression via promoter binding. As a result of the investigation described below, we hereafter refer to these proteins as Cas Protein Repressors (CasPRs), and assigned the identifier CasPRIB1 for AcrIB14, and CasPRIB2 for AcrIB12. We analyzed transcriptome-wide effects during expression of CasPRIB1 or CasPRIB2, which corroborated strong reductions in the levels of type I-B cas gene transcripts for both proteins, with minimal or inconsistent effects on other transcripts (**Fig. 1F**). As seen with RT-qPCR, the levels of cas6 mRNA were the least affected by either CasPR.

### CapR and AP2 domains are required for CasPR Acr activity

An HHPred analysis of CasPRIB1 and CasPRIB2 revealed that both proteins encode CapR and AP2 domains predicted to mediate DNA binding, raising the possibility that they transcriptionally regulate *cas* gene expression (**Fig. 2A**). CapR domains were discovered in phage-inducible satellite elements, where they mediate transcriptional repression of phage genes, and are also found in homing endonucleases within phage-borne group II introns^20,21^. AP2 domains can also be found alongside HNH nuclease domains in homing endonucleases, as well as in transcription factors in both bacteria and eukaryotes, with well-characterized representatives playing roles in plant development^22,23^. While CasPRIB1 contains a single copy of each domain, CasPRIB2 is substantially longer, harboring two N-terminal CapR domains, a central AP2 domain, and four C-terminal HTH domains (**Fig. 2A**). AlphaFold3 models of both CasPRs predict that a series of helical linkers connect these domains (**Fig. 2B-C**). Previous studies identified AP2 domain-containing genes in *acr* loci of *Listeria* and *Clostridiodes* prophages, however these genes did not mediate detectable inhibition against *L. monocytogenes* type II-A or *C. difficile* type I-B CRISPR-Cas systems, respectively^24,25^. Accordingly, it was hypothesized that DNA-binding by the AP2 domain may serve to regulate *acr* gene expression, similar to Acas. To investigate the necessity of CapR, AP2, and HTH domains for CasPR-mediated Acr activity, we engineered mutants lacking each domain and tested their ability to inhibit plasmid targeting by type I-B CRISPR (**Fig. 2D**). While deletions of the CapR or AP2 domains from either CasPR abolished inhibition of CRISPR interference, a CasPRIB2 mutant lacking all four HTH domains retained inhibitory activity. Previous studies have shown that CapR domains adopt a zinc finger-like fold, likely using four closely oriented cysteine residues to coordinate a Zn^2+^ ion necessary for interaction with DNA^20^. We performed sequence alignment of 16 CasPR homologs and identified universally conserved C33, C35, C37, C56, and C58 residues, which are found in close proximity to one another in the AF3 model (Fig. S2).

**Figure 2.**
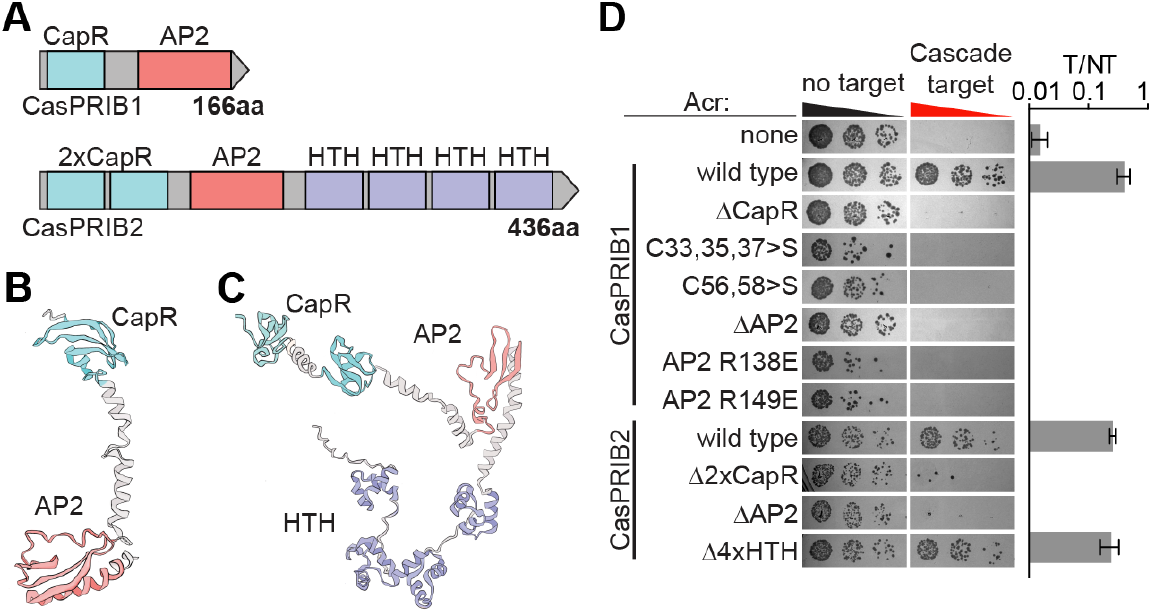
CapR and AP2 domains are required for CasPR activity. **(A)** Schematic of indicated CasPR proteins showing CapR, AP2, and HTH (helix-turn-helix) domains. **(B-C)** AlphaFold3 model of CasPRIBI (B) and CasPRIB2 **(C). (D)** Plasmid targeting assay testing inhibitory activity of the indicated CasPR proteins and mutant derivatives against the LS1 type l-B CRISPR-Cas system. T/NT, ratio of target to non-target transconjugants. Error bars indicate SEM.

Furthermore, we identified positively charged residues (R138 and R149) conserved within AP2 domains of *Listeria* CasPR homologs. We constructed CasPRIB1 alleles containing triple (C33/35/37S) or double (C56/58S) mutations to serine, and found that both lost Acr activity (**Fig. 2D**). We tested charge reversal mutations in the CasPRIB1 AP2 residues (R138E and R149E) and found that both abolished Acr activity. Together, these data show that CapR and AP2 domains are both necessary for CasPR activity while the HTH domains of CasPRIB2 are dispensable.

### CasPRs are abundant in Bacillota prophages

A stringent BLASTp search of the NCBI nr database for CasPRIB1 homologs yielded 4,542 unique hits across bacteriophages and diverse genera of phylum Bacillota, including *Listeria, Clostridioides, Streptococcus*, and *Enterococcus* (**Fig. 3A and S3A-B**), while other phyla largely lack high-confidence homologs. **Listeria** CasPR homologs form two distinct clades (A and B) with which CasPRIB1 and CasPRIB2 cluster, respectively. Manual inspection of selected leaves along the CasPR phylogenetic tree (**Fig. 3A and S3A**) indicated that many *caspr* genes are part of probable acr loci, located alongside homologs of previously identified *acr* genes residing within prophages. We tallied instances of *caspr* genes and 137 other defense inhibitor families in bacterial genomes, which revealed that CasPRs are the second most abundant inhibitor class in Bacillota, after AcrIIA21 (**Fig. S3C**). However, as AcrIIA21 is highly homologous to plasmid replication initiator proteins, we suspect that many AcrIIA21 homologs detected here do not play anti-CRISPR roles. We tabulated homologs of known *acr* genes located within 5 kb of a *caspr* gene in CasPR-containing bacterial genomes. This analysis revealed 9,118 clusters, with most containing putative type I-B (AcrIB8, IB5) and type II-A (AcrIIA1, IIA12) Acrs encoded near *caspr* genes (**Fig. S3D**). Finally, we searched for CasPR homologs within viral (IMGVR) and plasmid (IMGPR) sequence databases. While we uncovered predicted CasPRs within both classes, unique phage sequences harboring *caspr* genes (14,565 instances, 0.26% of sequences) outnumbered those in plasmids (324 instances, 0.046% of sequences) (**Fig. S3E**).

**Figure 3.**
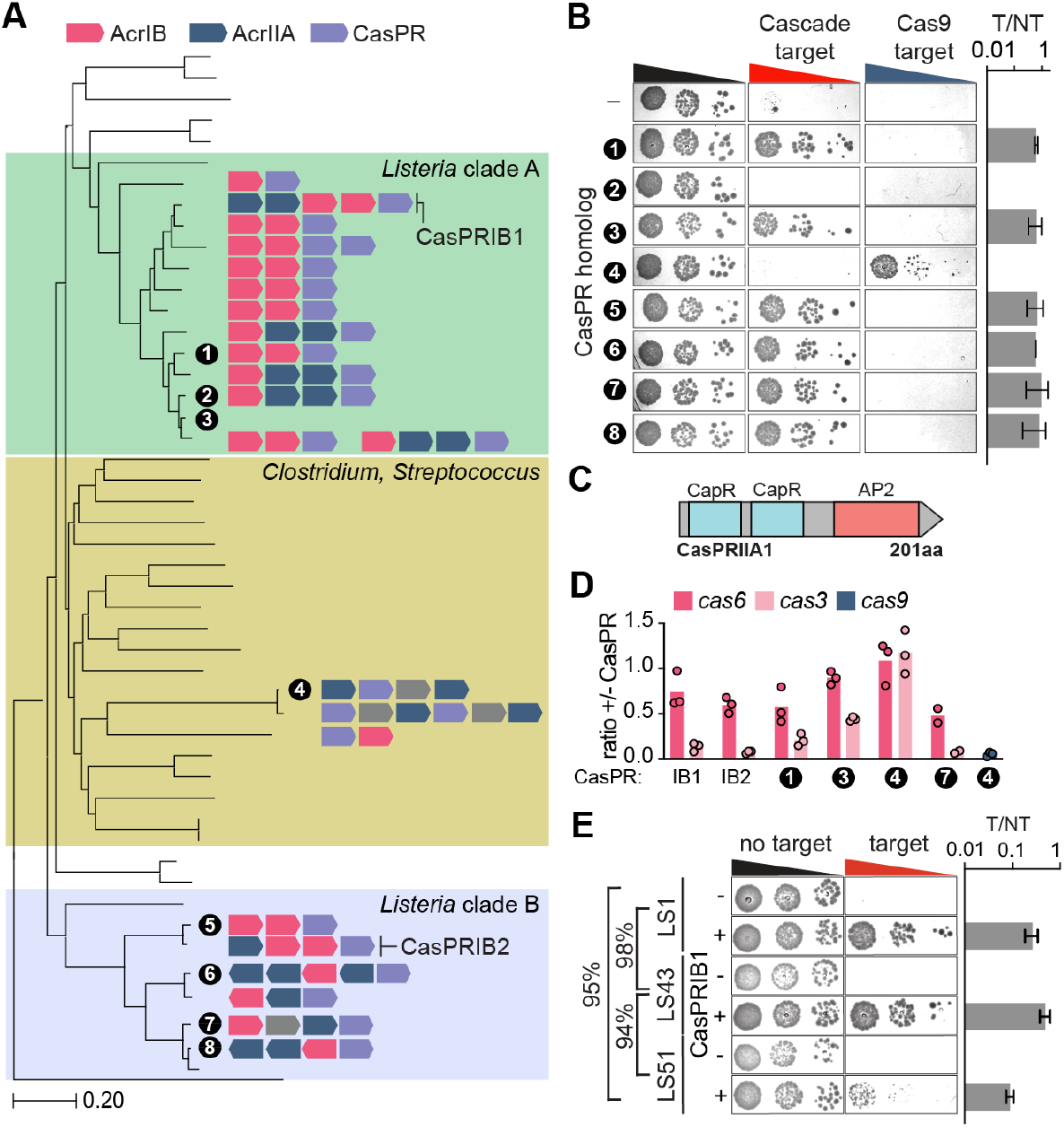
CasPR homologs inhibit diverse CRISPR types. **(A)** Phylogenetic tree representing a subset of Listeria, Streptococcus, and Clostridium spp. encoding casPR homologs. Acr clusters encoding casPR homologs (violet) for each leaf are depicted. Homologs tested for activity are labeled alongside their respective acr gene clusters. Type l-B acr homologs in pink, type ll-A in blue, gray genes indicate hypothetical proteins. See Fig. S3A for more detailed tree. **(B)** Plasmid targeting assays for homologs tested against type l-B (Casacde) and type ll-A(Cas9) CRISPR. T/NT, ratio of target to non-target transconjugants for Cascade target. Error bars are SEM. **(C)** Domain architecture of CasPRIIAI inhibitor (homo-log 4) from *Streptococcus dysgalactiae*. **(D)** RT-qPCR measuring transcript abundance of *cascade* (*cas6* and *cas3*) or *cas9* in the presence of CasPR homologs, as in Fig. 1D. **(E)** Plasmid targeting assays testing activity of CasPRIBI against diverse Cascade sequences encoded by 3 unique *L seeligeri* strains. Nucleotide sequence conservation across the cascade locus is depicted between strains. Error bars are SEM.

### Streptococcus dysgalactiae CasPRIIA1 inhibits Cas9

To investigate the generalizability of CasPR activity, we selected eight diverse CasPR homologs for experimental characterization from the phylogenetic tree in **Fig. 3A**, which range from 26% to 90% pairwise sequence identity and contain representatives from both Listeria clades (**Fig. S3F**). We cloned each into a plasmid under the control of a constitutive promoter, and tested their ability to inhibit plasmid targeting by the *L. seeligeri* type I-B CRISPR-Cas system (**Fig. 3B**). Six out of eight tested homologs inhibited interference by Cascade, with potency similar to that observed with CasPRIB1 and CasPRIB2. As type II-A Acrs are often encoded near *caspr* genes, we also tested the ability of each homolog to inhibit a type II-A CRISPR-Cas system from *L. seeligeri*, which we cloned into an integrating plasmid under the control of its native promoter (**Fig. 3B**). While none of the six CasPR homologs active against type I-B affected type II-A CRISPR interference, one of the two remaining homologs inhibited Cas9-mediated plasmid targeting (**Fig. 3B and S3G**), indicating that CasPR family members have evolved to antagonize completely unrelated CRISPR-Cas systems. This CasPR (here called CasPRIIA1) is the most divergent homolog in the tested set (**Fig. S3F**), is encoded by a prophage within *Streptococcus dysgalactiae* strain Kdys0611, alongside homologs of two other type II-A Acrs, and like CasPRIB2, contains two CapR domains (**Fig. 3A,C and S3A**). CasPRIIA1 also inhibited targeting by *Streptococcus pyogenes* Cas9 (which is 53% identical to LseCas9) in the same assay (**Fig. S3G**). Next, we performed RT-qPCR to investigate whether a selected subset of active CasPR homologs decreased *cas* transcript abundance similarly to CasPRIB1 and CasPRIB2 (**Fig. 3D**). CasPR homologs 1, 3, and 7 substantially decreased *cas3* transcripts, with limited or no effects on *cas6*. As expected CasPRIIA1 (homolog 4) had no effect on *cas6* or *cas3* transcript abundance, but strongly reduced *Lsecas9* mRNA levels. Taken together, these results indicate that diverse CasPRs inhibit CRISPR interference by reducing *cas* transcript levels.

Next, we explored the inhibitory range of an individual CasPR protein. We cloned type I-B CRISPR systems from *L. seeligeri* strains LS43 and LS51 into an integrating plasmid, equipped each with a plasmid-targeting spacer, and confirmed that both restricted a targeted plasmid (**Fig. 3E**). We then tested their susceptibility to inhibition by CasPRIB1. The LS43 and LS51 cas gene sequences are 98% and 95% identical to the previously tested LS1 CRISPR system at the nucleotide level, respectively, including nonsynonymous variation in all cas gene sequences. We observed that CasPRIB1 inhibited interference by the LS43 and LS1 type I-B systems with similar potency. However, CasPRIB1 only partially enabled tolerance of a plasmid targeted by the LS51 type I-B system, resulting in fewer and smaller target plasmid-containing transconjugant colonies than seen during inhibition of the other two systems (**Fig. 3E**). Thus, natural variations in type I-B CRISPR system sequence affect susceptibility to CasPRIB1.

### CasPRs bind site-specific motifs within cas genes

We hypothesized that CasPRs inhibit synthesis of CRISPR-Cas machinery by binding to DNA within their respective CRISPR-Cas loci. Accordingly, we performed ChIP-seq using six functionally tagged CasPR-his6 alleles and plotted binding enrichment across the genome compared to an empty vector control strain lacking Acrs (**Fig. 4A-B and S4**). Strikingly, for each CasPR active against type I-B CRISPR, we observed a sharp peak of CasPR occupancy within the *cas8b* coding sequence toward the 5’ end of the gene. (**Fig. 4A and S4**). That the putative binding site for each type I-B CasPR lies within *cas8b* explains our observations that CasPRs have limited effects on transcripts of *cas6* (the first gene in the operon, which is located just upstream of the site) but strong effects on downstream genes (**Fig. 1E and 3D**). In contrast to type I-B CasPRs, CasPRIIA1 occupancy was not enriched anywhere within the type I-B CRISPR-Cas locus, but was strongly enriched near the 5’ end of the *Lsecas9* coding sequence (**Fig. 4A-B**).

**Figure 4.**
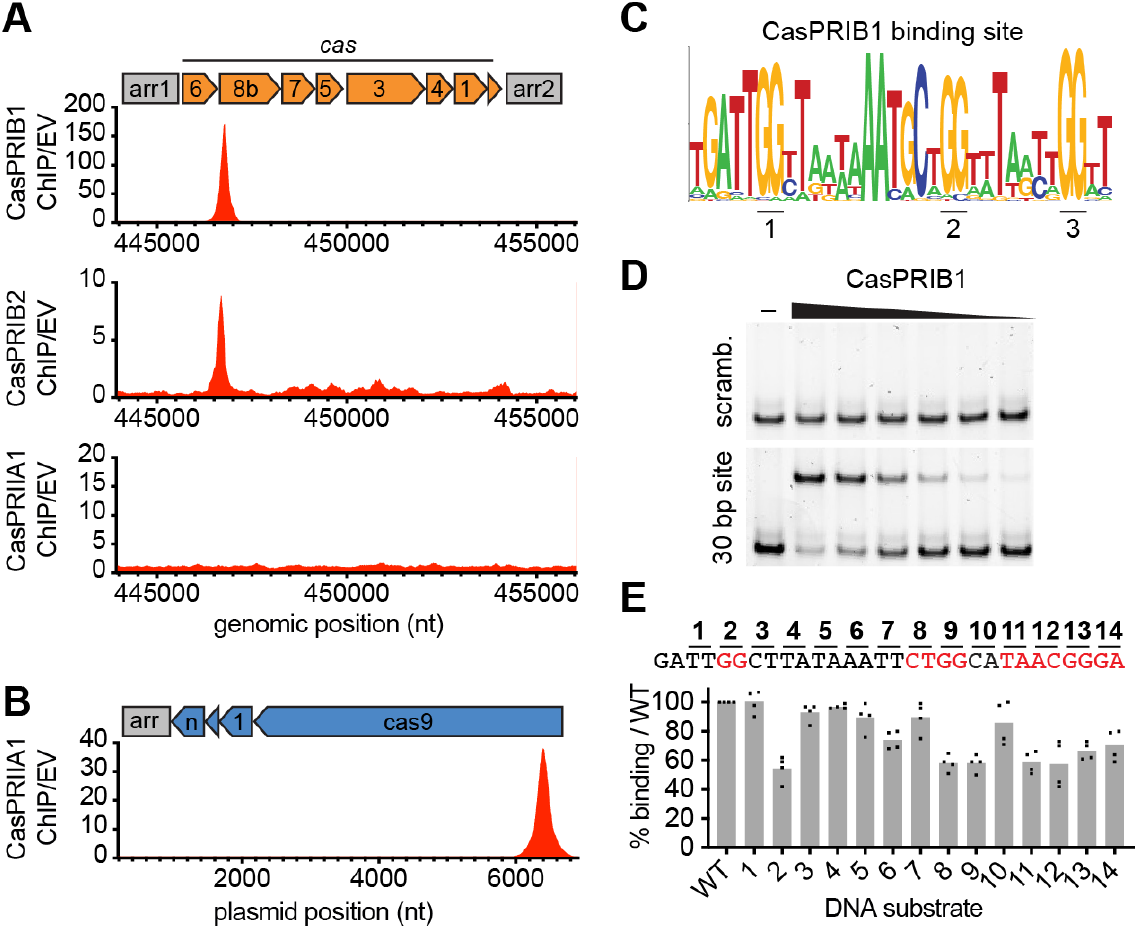
Site-specific binding of CasPRs to cas genes. **(A)** ChlP-seq profiles of indicated his6-tagged CasPRs, plotted along the LS1 type l-B CRISPR-Cas locus. Plot represents the ratio of reads in CasPR ChIP samples to those in empty vector (EV) control lacking CasPRs. ChIP signal was normalized to reads from a total input DNA sample. **(B)** As in (A), but plotting the CasPRIIAI ChIP profile along the type ll-A CRISPR-Cas locus from L. seeligeri. **(C).** Weblogo assembled from alignment of 250 unique cas8b gene sequences, representing region bound by CasPRIBI. **(D)** EMSA with 50 nM of indicated 60 bp DNA substrate and 0, 250, 200, 150, 100, 75, 50 nM CasPRIBI, resolved on native acrylamide gels. **(E)** Sequence in the cas8b gene determined critical for CasPRIBI binding in EMSAs (See Fig. S5). Binding measurements with 150 nM CasPRIBI and 50 nM indicated substrates containing 2 bp mutations with respect to the WT 30 bp CasPRIBI binding site. Percent binding relative to WT substrate shown for 4 biological replicates. Red nts exhibit diminished binding when mutated.

Elsewhere in the genome, diverse CasPRs exhibited unique but reproducible binding profiles, which ranged from four to over one hundred putative binding sites (**Fig. S4**). Despite this, RNA-seq analysis during CasPRIB1 and CasPRIB2 expression showed that *cas* genes are the primary targets of silencing (**Fig. 1F**). Each ChIP peak exceeded 200 bp in width, complicating the identification of a consensus binding motif. Therefore, we refined the CasPRIB1 binding site by conducting gel shift assays with candidate DNA targets. We expressed CasPRIB1 in *E. coli* and purified it to near homogeneity (**Fig. S5A**). When analyzed via gel filtration, CasPRIB1 eluted at a volume consistent with a monomeric state (**Fig. S5B**). Notably, we found that CasPRIB1 bound much more efficiently to a heparin column when purified from cultures supplemented with Zn-acetate, which along with the identification of functionally important conserved cysteines (**Fig. 2D**) suggest that coordination of a Zn^2+^ ion may be important for its DNA-binding activity. Accordingly, we established in vitro DNA binding assays using CasPRIB1 purified from Zn-treated cultures. We first observed specific CasPRIB1 binding to a 250 bp substrate containing *cas8b* DNA spanning the ChIP peak, and subsequent truncations of this fragment narrowed the binding activity to a 126 bp segment (**Fig. S5C**). To refine the site further, we analyzed twelve 60-bp tiled substrates, each overlapping by 5 bp, that collectively covered the 120-bp interval (**Fig. S5D-E**). This analysis identified the 5’ and 3’ boundaries of a 30-bp sequence required for CasPRIB1 binding (**Fig. 4C and S5D-F**). Using a DNA substrate containing this sequence (flanked by 15 bp on either side with non-specific DNA), we confirmed that CasPRIB1 specifically bound this 30 bp target sequence (**Fig. 4D**). AP2 domain-containing proteins often bind to short G/C-rich motifs^26,27^, and alignment of 250 *cas8* nucleotide sequences revealed the presence of 3 guanine dinucleotides among the highly conserved nucleotides within the 30 bp region (**Fig. 4C**). We introduced a series of 2 basepair non-overlapping mutations walking the length of the target and quantitated CasPRIB1 binding to each mutant substrate (**Fig. 4E**). Substrates containing mutations in any of the three GG pairs, or selected nucleotides clustered toward the 3’ end of the site, exhibited substantial reductions in CasPRIB1 binding. We tested CasPRIB1 binding to seven diverse 30 bp sequences from natural *cas8b* genes which vary along the region. CasPRIB1 strongly bound all but three of these substrates, each of which varied at position 18 (T>G or T>C) and/or position 24 (A>G or A>T) (**Fig. S5F**), suggesting roles for these nucleotides in CasPRIB1 binding. Finally, the CasPRIB1 binding site in the partially CasPR-resistant *L. seeligeri* str. LS51 *cas8b* gene differs at 10 out of 30 positions (including position 24) from its CasPR-sensitive homologs in strains LS1 and LS43, which share 100% identity with one another at this site (**Fig. 3E and S5F**). Together, these data indicate that CasPRs bind to specific motifs within *cas* gene DNA.

### CasPRIB1 DNA binding is necessary and sufficient to inhibit transcription

Next, we investigated the relationship between CasPRIB1 binding to its recognition site within *cas8b* and inhibition of CRISPR-Cas function. The 30 bp binding site lies within the *cas8b* coding sequence, and the three GG dinucleotides necessary for efficient CasPRIB1 binding are part of the W12 (TGG), G17 (GGC), and G20 (GGA) codons. While evolutionary pressure on the host to resist CasPR-mediated silencing could drive mutations in the binding site, the mutation of any individual GG pair in this context necessitates a nonsynonymous mutation that might affect CRISPR function. We explored this concept by using a plasmid-targeting assay to test the activity and CasPR-sensitivity of *cas8b* alleles mutated in each of the three GG pairs, and a recoded allele that left all three GG pairs intact (**Fig. 5A**). Mutation of the first and third GG (W12A and G20S) abolished CRISPR function, suggesting these nucleotides may be mutationally constrained. In contrast, mutation of the second GG (G17Y), or recoding of the site, did not affect CRISPR function. Both the G17Y and recoded alleles became resistant to CasPRIB1, indicating that CasPR binding of its recognition site is required for inhibition of CRISPR immunity (**Fig. 5A**). That the recoded site lost susceptibility to inhibition supports the notion that other nucleotides in addition to the three GG pairs contribute to CasPRIB1 binding.

**Figure 5.**
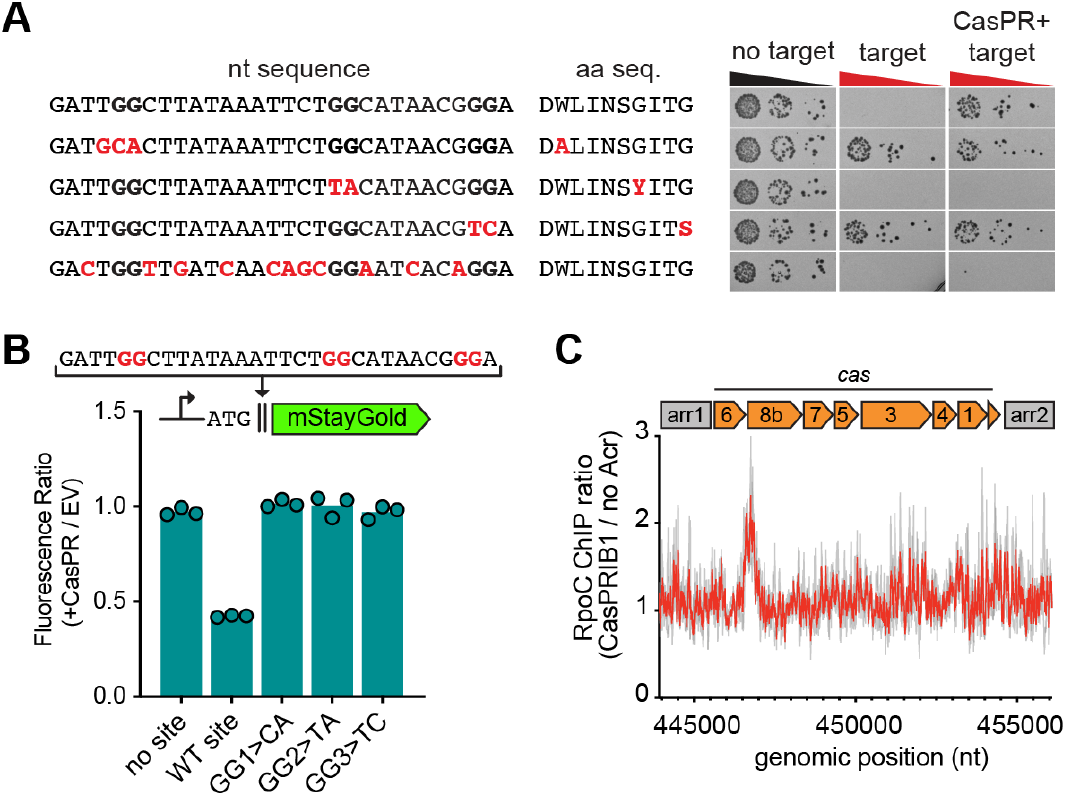
CasPR binding to target DNA is necessary and sufficient to block transcription, **(a)** Plasmid targeting assays testing effects of the indicated (red) cas8b mutations on CRISPR function and CasPRIBI sensitivity. Changes to amino acid sequence resulting from mutations are shown in red. Donor plasmids with or without a type l-B CRISPR target were introduced into strains harboring the indicated mutation in the presence and absence of a CasPRIBI-expressing plasmid. **(B)** Fluorescent reporter assay testing sufficiency of CasPRIBI DNA binding for transcriptional inhibition. The indicated CasPRIBI binding sites or mutant derivatives were inserted in-frame ahead of an mStayGold reporter, downstream of start codon. Fluorescence of each reporter variant was measured in the presence and absence of CasPRIBI. Plot reports ratio of fluorescence in presence of CasPRIBI to that without CasPR. **(C)** RNAP enrichment at CasPR binding site in presence of CasPR expression. Profile represents ratio of RpoC-his10 chip signal in presence of CasPRIBI to that without, plotted along the L. seeligeri type l-B CRISPR-Cas locus. Gray traces indicate two biological replicates,

We tested whether CasPR binding to DNA is sufficient to block transcription of downstream genes. We inserted the 30 bp CasPRIB1 binding site in-frame ahead of a chromosomal mStayGold fluorescent reporter immediately following its start codon (**Fig. 5B**). We also generated reporter variants harboring mutations in each individual GG dinucleotide. We introduced a plasmid expressing CasPRIB1, and measured reporter fluorescence versus empty vector (no CasPR) controls. We saw that CasPRIB1 reproducibly drove a 2-fold reduction in fluorescence, specifically in the presence of its binding site (**Fig. 5B**). Furthermore, mutation of any guanine pair abolished the repressive effect. These results indicate that CasPRIB1 binding to DNA is sufficient to repress transcription of downstream genes and corroborate the roles of all three GGs in CasPRIB1 recruitment in vivo.

Binding within a coding sequence is an atypical mode of gene repression in prokaryotes. Our RT-qPCR and RNA-seq data indicate that CasPRIB1 affects expression of genes downstream of its binding site (*cas8b, cas7, cas5, cas4, cas3*) but not upstream (*cas6*) (**Fig. 1E-F**). Hence we hypothesized that DNA-bound CasPRIB1 may present a transcriptional roadblock to elongating RNA polymerase. To investigate this, we tagged the β’ subunit of RNA polymerase at its native locus (*rpoC-his10*) and performed ChIP-seq to measure RNAP occupancy on DNA in the presence and absence of CasPRIB1 (**Fig. 5C**). We found that expression of CasPRIB1 resulted in approximately 2-fold enrichment of RNAP at the CasPR binding site compared to a no Acr control, indicating that CasPRIB1 causes accumulation or pausing of RNAP during transcription.

### Phage-encoded CasPRIB1 inhibits type I-B CRISPR during lysogeny

Finally, we investigated the effects of a phage-encoded CasPR on CRISPR immunity during infection. The temperate phage ϕEGDe naturally encodes three type II-A Acrs alongside CasPRIB1 (**Fig. 6A**). To test whether CasPRIB1 inhibits Cascade interference during phage infection, we performed plaque assays using lawns of *L. eeligeri* LS1 expressing type I-B CRISPR equipped with spacers targeting ϕEGDe, infected with either WT ϕEGDe or a mutant containing an in-frame deletion of the *casprIB1* gene (**Fig. 6B**). We found that both phages were susceptible to CRISPR targeting during lytic infection, suggesting that ϕEGDe *casprIB1* is either not expressed or incapable of neutralizing pre-existing Cascade targeting complexes under these conditions. In contrast to the circumstances of lytic infection, we hypothesized that prophage-expressed *casprIB1* might lower the steady-state levels of CRISPR-Cas machinery. Using both WT and *ΔcasprIB1* mutant phages, we lysogenized a strain of LS1 harboring type I-B CRISPR armed with a plasmid-targeting spacer. When we performed plasmid-targeting assays with these lysogens, WT ϕEGDe, but not ϕEGDe *ΔcasprIB1*, inhibited CRISPR immunity (**Fig. 6C**). Thus, ϕEGDe *casprIB1* is expressed and functional during lysogeny.

**Figure 6.**
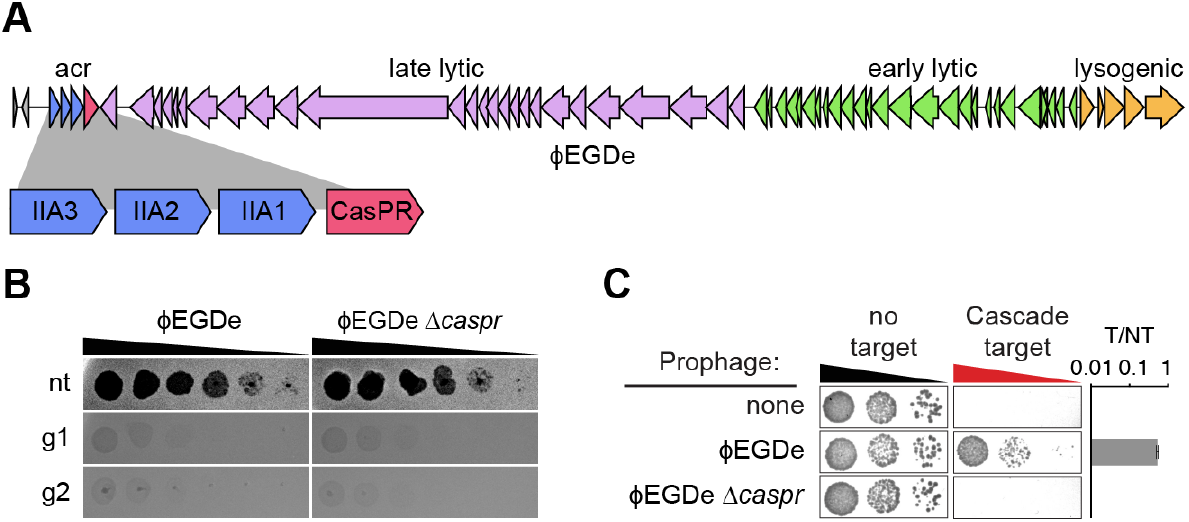
CasPRIBI inhibits CRISPR interference during lysogeny. **(A)** Schematic of the ϕ EGDe phage genome, highlighting acr locus which contains casprIBI homolog. **(B)** Plaque assays with ten-fold serial dilutions of the indicated phage plated on lawns of L. seeligeri with non-targeting (nt) or two different targeting type l-B gRNAs (g1 and g2). **(C)** Plasmid targeting assay testing inhibitory activity of the indicated prophage and mutant derivative on the LS1 type l-B CRISPR-Cas system. T/NT, ratio of target to non-target transconjugants. Error bars indicate SEM.

## Discussion

Here we characterized CasPRs as DNA-binding anti-CRISPRs widespread in Bacillota phages, that transcriptionally silence diverse CRISPR-Cas systems by recognition of specific sites within *cas* genes. While we identified eight unique CasPRs that silence type I-B and one that silences type II-A systems, we hypothesize that additional CasPRs exist with even more diverse specificity. Recognition of DNA targets rather than Cas proteins themselves may enable evolutionary flexibility for CasPRs to rapidly adapt to silence other defenses or competitors. Conceivably, CasPRs could evolve to silence other CRISPR systems, non-CRISPR defenses, and perhaps the lytic genes of competing phages.

Additional study is needed to define the relationship between the CapR and AP2 domain structures and the target sequences they recognize within *cas* genes. For example, some CasPRs contain multiple CapR domains, which may confer additional stringency to their binding site, while CasPRs with a single CapR domain may be more relaxed. We also identified CasPRs with additional HTH domains dispensable for Acr activity, and hypothesize that these domains may serve an autoregulatory function, similar to an Aca. Rather than eliciting repression by binding to *cas* gene promoters, our data indicate that diverse CasPRs bind within coding sequences, which are unusual sites for prokaryotic transcriptional regulation. One possibility driving the targeting of coding sites is that they may be more mutationally constrained than noncoding ones, preventing evolutionary escape from repression, as we observed for key nucleotides in the CasPRIB1 binding site within *cas8b*.

Recently, a parallel mechanism was reported to mediate *cas* gene repression by some Pseudomonas phages and plasmids^28^. A subset of CRISPR-Cas systems avoid autoimmunity by producing crRNA-like RNAs with imperfect homology to the *cas* gene promoter, which direct Cas nucleases to autorepress their own synthesis^29,30^. Li and colleagues discovered viral homologs of these RNAs (called cracrs) that exploit this regulatory circuit to indirectly enforce *cas* repression. Thus, diverse invading entities have evolved independent mechanisms to transcriptionally regulate host immune systems. Repression by cracrs is likely limited to CRISPR systems that utilize regulatory RNAs and depends upon host Cas nucleases. In contrast, CasPRs directly regulate *cas* gene expression without relying on host proteins.

What roles do CasPRs play in the phage lifecycle? We determined that CasPRIB1 from phage ϕEGDe inhibits type I-B CRISPR immunity during lysogeny, but is insufficient to protect the phage from CRISPR interference during lytic infection. We suspect that neutralization of fully formed interference complexes in the infected cell requires a more rapid mechanism than transcriptional repression. It is worth noting that most CasPRs are encoded alongside other Acrs, including those targeting the same CRISPR type as the CasPR. Therefore, during infection conventional Acrs could bind and inactivate pre-existing CRISPR machinery while CasPRs block its replenishment. Similarly, prophages could benefit from the combined actions of CasPRs that reduce Cas protein synthesis with conventional Acrs that inactivate the few interference complexes that are nonetheless produced.

## Supporting information

Table S1. Strains Plasmids and Oligos used in this study

## Acknowledgements

We thank Barry Stoddard, Peter Fineran, Maximilian Feussner, Nils Birkholz, and all members of the Meeske Lab for helpful discussion. We thank Joseph Bondy-Denomy and Michelle Grunberg for pilot experiments with a putative *Pseudomonas* CasPR. SigA antibody was a kind gift of David Rudner. Work in the Meeske lab is supported by NIGMS (R35GM142460) and NSF (FAIN2235762). This material is based upon work supported by the Defense Advanced Research Projects Agency (DARPA) under Agreement No. HR00112590142. AJM is a Rita Allen Foundation Scholar. EMS is supported by T32GM136534 and the NSF Graduate Research Fellowship.

## Author Contributions

The study was conceived by EMS and AJM. EMS, LO, VMH, and AJM performed all in vivo experiments and associated analyses. SN, JP, and BKK purified CasPR proteins and performed in vitro DNA-binding assays. EMS, BKK, and AJM wrote the paper. All authors contributed to editing the paper.

## Declaration of Interests

AJM is a co-founder and advisor of Profluent Bio. The other authors declare no competing interests.

## Materials and Methods

### Bacterial strains and growth conditions

*Listeria seeligeri* strains were cultured with Brain Heart Infusion (BHI) media at 30°C and supplemented with one or more antibiotics at the following working concentrations: nalidixic acid (50 μg/mL), chloramphenicol (10 μg/mL), erythromycin (1 μg/mL), or kanamycin (50 μg/mL).

*Escherichia coli* strains utilized for cloning and conjugative plasmid transfer were cultured with Lysogeny Broth (LB) media at 37°C and supplemented with antibiotic at the following working concentrations: ampicillin (100 μg/mL), chloramphenicol (25 μg/mL), kanamycin (50 μg/mL).

Shuttle vectors utilized in conjugative transfer were purified from Turbo Competent *E. coli* (New England Biolabs) and transformed into the *E. coli* conjugative donor strains SM10 λpir or S17 λpir. For a list of strains used in this study, see Table S1.

### Plasmid construction and preparation

Genetic constructs used for expression in *Listeria seeligeri* were cloned into one of two shuttle vector backbones described below using Gibson Assembly of digested plasmid and PCR amplified inserts. All plasmids used in this study and their construction details can be found in Table S1.

pPL2e: A single-copy plasmid that integrates into the tRNA^Arg^ locus within the *L. seeligeri* chromosome. Confers chloramphenicol resistance in *E. coli* and erythromycin resistance in *L. eeliger*i^31^.

pAM326: A multi-copy plasmid conferring kanamycin resistance in *E. coli* and *L. seeligeri*^32^.

CRISPR-Cas mutant constructs were cloned using KLD site-directed mutagenesis methods with primers containing mutant binding sequences to amplify a type I-B CRISPR-Cas expression plasmid^11^. Anti-CRISPR constructs and associated mutants were cloned into NcoI/EagI digested pAM551 backbones originally derived from pAM326 but including an aTc-inducible Ptet promoter.

### *E. oli – L. seeligeri* conjugation

Plasmids were introduced to strains of *L. seeligeri* using *E. coli* conjugative donor strains SM10 λpir or S17 λpir. 100 μL of donor *E. coli* and 100 μL of recipient *L. seeligeri* turbid cultures were diluted into 5 mL of BHI broth and concentrated on a 0.45 μM filter using vacuum filtration. Filters were placed cell side up on BHI agar plates supplemented with oxacillin (8 μg/mL for pPL2e- or pAM326-derived plasmids and 128 μg/mL for pAM8-derived plasmids) and incubated at 37°C for 4 hours. For strain construction, cells were streaked directly from the filter onto BHI agar plates containing appropriate antibiotics. When performing plasmid targeting assays to test CRISPR interference activity, filters were resuspended in 2 mL BHI broth, and 5 μL of ten-fold serial dilution of cell suspension was spotted onto BHI agar plates containing appropriate antibiotics, including nalidixic acid to kill donor *E. coli*. Plates were incubated for 48-72 h at 30°C.

### *L. eeligeri* strain and phage engineering

Chromosomal mutations in *L. seeligeri* were generated by allelic exchange as previously described^33^. Briefly, repair templates were prepared by cloning 1 kb fragments containing homology upstream and downstream flanking the mutation site into the suicide plasmid pAM215 (lacZ+ cmR). To generate *rpoC-his10*, the repair template pAM724 was conjugated into *L. eeligeri* str. LS1. To generate ϕEGDe *ΔcasprIB1*, the repair template pES103 was conjugated into LS1 lysogenized with ϕEGDe. Transconjugants were selected on BHI containing nalidixic acid and chloramphenicol, then passaged three times in the absence of selection by growing to saturation in BHI broth at 30°C and diluting 1:1,000 in fresh media. Cells were diluted 106-fold and plated on BHI agar plates supplemented with X-gal (100 μg/mL). After 3 days incubation at 30°C, white colonies were selected and analyzed for the desired mutation by colony PCR and Sanger sequencing. To generate a stock of ϕEGDe *ΔcasprIB1*, the resulting mutant lysogen was cultured to saturation in 5 mL BHI media, then subcultured in 5 mL BHI supplemented with 2 μg/mL mitomycin C overnight. The lysed culture was filtered using a 0.45 μm syringe filter, and plated with a top agar lawn of LS1 overnight. A single plaque was picked and expanded for stock generation, and confirmed by PCR to contain the *casprIB1* deletion.

### Western blots

*Listeria seeligeri* strains harboring His6-tagged proteins were grown in 5 mL cultures at 30°C in BHI + antibiotic to saturation. 1mL of overnight culture was pelleted via centrifugation and resuspended in 50 μL lysis buffer (20 mM Tris pH 7.5, 1 mM EDTA, 10 mM MgCl2, 2 mg/mL lysozyme, 0.4 mg/ mL DNAse I, 0.2 μg/mL RNAse A, and 1 uM PMSF). Cell suspensions were incubated in a 37°C water bath for 30 min and subsequently diluted in 50 μL of 2x Laemmli buffer (containing 4% SDS and 10% beta-mercaptoethanol) then boiled at 95°C for 5 min. Samples were then loaded into a 4-20% precast mini protean TGX gel running cassette (Bio-Rad) and run in 1x SDS-PAGE running buffer. Protein bands were transferred to a PVDF membrane for 1 hour and washed with PBST (1X PBS + 0.05% Tween-20) before incubating with primary anti-His antibody (Genscript) (1:4,000 dilution in 3% bovine serum albumin and 0.2% sodium azide) or anti-sigA antibody (gift from David Rudner) (1:10,000 dilution in 3% bovine serum albumin and 0.2% sodium azide) overnight. The following day, the membrane was washed with PBST and incubated with 2 μL HRP-conjugated mouse (His) or rabbit (sigA) secondary antibody (1:3,000 diluted in PBST) in 5% milk for 1h. Membrane was washed with PBST and HRP detection reagents (ThermoFisher) were utilized according to the manufacturer’s instructions to visualize bands.

### RT-qPCR

#### RNA extraction and purification

Strains of L. seeligeri were grown in 5 mL cultures at 30°C in BHI containing appropriate antibiotics to midexponential phase (OD600 = 0.3-0.8). Cultures were pelleted via centrifugation and pellets were resuspended in 100 μL of 1X PBS with 2 mg/mL lysozyme and incubated for 5 min in a 37°C water bath. After incubation, 10 μL of 10% sarkosyl was added to each sample and resuspended before adding 300μL TRI reagent. Samples were vortexed and 400μL of 100% ethanol was added. Each sample solution was then purified using the Zymo Direct-zol RNA Miniprep Kit to the manufacturer’s instructions. RNA-extraction samples were treated with DNAse and column purified using the Zymo RNA Clean & Concentrator kit per the manufacturer’s instructions.

#### qPCR

PCR was performed in 384-well plates in 10 μL reaction volumes using the NEB Luna Universal One-Step RT-qPCR Kit. PCR reactions utilized primer pairs that recognized individual type I-B and type II-A *cas* genes. Measurement of housekeeping gene ssbA transcripts was used to normalize total transcript abundance across samples. Relative cas gene expression for each sample was calculated as 2^ΔCt(*cas - ssbA*).

### RNA-seq

RNA was purified as described above from strains expressing a chromosomally integrated Ptet-driven type I-B CRISPR-Cas locus and either empty vector, or plasmid-borne copies of CasPRIB1 or CasPRIB2, in biological triplicate. RNA-seq library preparation and sequencing was performed by Genewiz, including rRNA depletion, DNase treatment, and 2×150 paired- end sequencing on an Illumina NextSeq platform. Reads were mapped to the *L. seeligeri* LS1 genome using Bowtie2^34^ with settings -I 10 and -X 700. Reads per kilobase values for each gene in the LS1 genome were calculated from the resulting BAM files using Subread featureCounts^35^, dividing gene read count values by gene length and applying a scale factor to normalize each library to 10M total mapped reads.

### ChIP-seq

#### Crosslinking and Immunoprecipitation

Strains of *L. seeligeri* expressing CRISPR-Cas systems and CasPR-His6 or RpoC-His10 proteins were grown in 50 mL cultures at 30°C in BHI + antibiotic to mid-exponential phase (OD 0.3-0.8). Cultures were crosslinked with 1% formaldehyde in 10 mM sodium phosphate at room temperature for 10 min before quenching with 2.5 mL of 2.5 M glycine for 5 min. Cultures were pelleted via centrifugation at 4°C and washed 3 times by resuspension in 10 mL ice cold 1X PBS. After washes, pellets were resuspended in 1.5 mL lysis buffer (50 mM HEPES pH 7.0, 150 mM NaCl, 5 mM MgCl2, 10 mM imidazole, 1 mg/mL lysozyme, 1 mM PMSF, 5% glycerol), and incubated at 37°C for 15 min. Samples were sonicated (Qsonica Q500) on ice using 15% amplitude in 10 s intervals with a 50% duty cycle for 10 minutes of processing time, which resulted in DNA fragments averaging 500 bp.

Processed lysates were centrifuged for 10 minutes at 4°C. Here, 50 μL of clarified lysates were reserved as “total DNA” matched pairs for each sample. Remaining supernatants were applied to 100 μL of equilibrated Ni-NTA agarose resin (ThermoFisher) and incubated at 4°C with rotation for two hours. Then, mixtures were centrifuged at 4°C and 2000 rpm for 1 min to pellet beads. Beads were washed with 1 mL of cold wash buffer (20 mM HEPES pH 7.0, 150 mM NaCl, 5 mM MgCl2, 10 mM imidazole, 5% glycerol) and further incubated at 4°C for 5 min. This was repeated twice for three total washes.After the final wash, 50μL elution buffer (20 mM HEPES pH 7.0, 150 mM NaCl, 5 mM MgCl2, 250 mM imidazole, 5% glycerol) was added to each bead pellet. Samples were incubated at RT for 5 minutes and centrifuged at 4°C and 2000 rpm for 1 min before carefully harvesting the IP supernatant. Total DNA and IP samples were incubated at 60°C overnight to reverse crosslinks.

#### DNA purification

After overnight incubation, 450 μL of wash buffer was added to each sample and samples were treated with RNase A (0.2 mg/mL) at 37°C for 1 h and subsequently with Proteinase K (0.2 mg/mL) at 37°C for 30 min. Next, an equal volume of phenol/chloroform (1:1) was added to each sample and vortexed before centrifuging samples at RT for 10 min at 13,000 rpm.

The aqueous phase of each sample was then harvested and combined with 50 μL sodium acetate. DNA was precipitated with two volumes of cold ethanol and pelleted by centrifugation (15,000 rpm) at 4°C for 15min. Pellets were washed two times with 70% ethanol before resuspending IP samples in 10 μL water and total DNA samples in 50 μL water.

#### Library Preparation and Sequencing

DNA fragments for each sample were barcoded and prepared for sequencing using the Illumina TruSeq DNA Library Preparation kit according to the manufacturer’s instructions. Purification steps were performed using Ampure beads and samples were pooled equally by weight to form final libraries. CasPR ChIP-seq libraries were sequenced by Azenta on the Illumina NextSeq platform (2×150 bp reads). RpoC-his10 libraries were sequenced on the Illumina MiSeq i100 platform (2×50 bp reads).

#### Data Analysis

Reads were mapped to the *L. seeligeri* genome using Bowtie2^34^ with settings -I 10 and -X 700. Coverage across the genome was calculated from the resulting BAM files using bamCoverage^36^ with a bin size of 1 nucleotide, and coverage was normalized to 10M reads per sample. Coverage at each nucleotide position in the resulting wig files was normalized twice: first the values for each IP sample were divided by the values in their corresponding total DNA sample, then the normalized coverage values in samples expressing CasPRs (or RpoC-his10) were divided by an empty vector control.

### Fluorescent reporter assays

A 30 bp CasPRIB1 binding site (or mutant derivative) was cloned in-frame ahead of the mStayGold fluorescent reporter gene, just after the stop codon. Reporter expression was driven by the constitutive promoter Phyper and was delivered to *L. seeligeri* using the integrating vector pPL2e. CasPR-expressing plasmids or empty vector controls were introduced into each reporter strain via a second round of conjugation. 1 mL of exponentially growing *L. seeligeri* cultures harboring each reporter was harvested at OD600 = 0.6 and pelleted by centrifugation. Cell pellets were each washed four times with 1 mL PBS, and resuspended in a final volume of 1 mL PBS. 200 μL cell suspension was aliquoted into black clear-bottom 96-well plates for analysis. Fluorescence was measured using a BMG Labtech CLARIOstar Plus plate reader, with the following settings: Excitation wavelength 470-15 nm, Dichroic auto 491.2 nm, Emission wavelength 515-20 nm, Gain adjustment 90%. OD600 was measured from the same samples, and these values were used to normalize fluorescence readings.

### CasPR homolog identification

CasPRIB1 homologs were identified by BLASTp search of the NCBI nr database using CasPRIB1 from *L. seeligeri* str. RR4 (accession: QDA75636.1) as query, with an E-value cutoff of 1E-4 and minimum query coverage of 70%. Taxonomy information was collected from all hits. 4,452 retrieved CasPR FASTA sequences were extracted and clustered using the T-Coffee seq_ reformat tool^37^ with a 90% sequence identity cutoff, resulting in 92 protein clusters. Phylogenetic tree was constructed from the cluster representatives using MEGAv12^38^ with the neighbor-joining method and 500 bootstrap iterations. To search for *caspr* genes within *acr* gene clusters, a BLAST database was assembled from 18,754 CasPR-encoding genomes extracted from the first search. This database was searched for both *casprs* and representatives of 137 other counterdefense proteins using TBLASTN with E-value cutoff 1E-4. Regions containing both a *caspr* gene and non-CasPR *acr* gene located within a 5 kb range were tabulated. To compare CasPR abundance with that of other Acrs in bacterial genomes, the NCBI nt_prok database of prokaryotic genomes was searched for genes encoding homologs of CasPRIB1 and 137 representative Acr families by TBLASTN with E-value cutoff of 1E-4 and minimum query coverage of 75%. Taxonomy information was collected for genomes containing at least one hit (additional hits within a single genome were not considered) and tabulated by Acr class. To compare CasPR content of phages and plasmids, the IMGVRv4^39^ and IMGPRv1^40^ databases were searched using CasPRIB1 as a query with E-value cutoff of 1E-4 and minimum query coverage of 75%. Phage and plasmid sequences encoding at least one CasPR were tallied.

### CasPRIB1 expression and purification

#### CasPRIB1 Expression

Codon-optimized CasPRIB1 for expression in *E. coli* was subcloned into pET21d+ without any additional affinity tags and transformed into BL21(DE3)pLysS bacteria.

Inductions were carried out by one of three methods:

LB media (IPTG): 20-50 mL overnight cultures started from single colonies were grown in LB supplemented with 100 μg/mL ampicillin. The next morning, 10 mL of culture was transferred into 2.8-L Fernbach flasks containing 1 L of pre-warmed LB with ampicillin (100 μg/mL) and incubated at 37°C with shaking at 205 rpm. When cultures reached an OD_600_ of 0.6, expression was induced by adding 0.5 mM IPTG. Cultures were then shifted to 18°C and shaken for 18–22 hours. Cells were harvested by centrifugation at 7,800 × g for 10 min at 4°C, resuspended in Buffer A (25 mM Tris–HCl, pH 7.5; 200 mM NaCl), transferred to 50 mL Falcon tubes, and spun again in a swinging bucket Eppendorf X rotor at 4,000 rpm for 15 min. Supernatants were discarded, and the resulting pellets were stored at −20°C.

LB media (IPTG) with Zn2+ addition: The same protocol above was used, except when the 1 L cultures reached an OD600 of 0.6, 0.2 mM Zn-acetate was added, incubated for 20 minutes, followed by addition of 0.5 mM IPTG.

Autoinduction^41^: ten colonies from a transformed plate were inoculated into 1 L of Autoinduction media (Including ZY media supplemented with trace metals) containing 100 μg/ mL ampicillin in 2.8 L flasks and shaken at 205 rpm for 9 h at 37°C followed by shaking for 24 hours at 18°C. Cells were harvested as described above.

#### CasPRIB1 purification

Frozen induction cell pellets were thawed and resuspended in ice cold 300 mM NaCl, 25 mM Tris pH 7.5 using ∼30 mL buffer per liter of induced culture, then lysed by sonication on ice. The lysate was clarified by centrifugation for 25 min at 18,000 rpm in a Sorvall SS-34 rotor, followed by filtration through a 5 μm syringe filter (Millipore). The resulting supernatant was loaded onto a 1 mL Heparin HP HiTrap column (GE Healthcare) pre-equilibrated in Buffer A using a peristaltic pump. The column was subsequently connected to an FPLC (AktaPrime, GE Healthcare) and eluted with a 30 mL linear gradient from Buffer A to Buffer B (25 mM Tris–HCl, pH 7.5; 1 M NaCl). Peak fractions were combined, concentrated to ≤2 mL, and further purified by size-exclusion chromatography (SEC) on a HiLoad 16/60 Superdex 200 prep grade column (Millipore Sigma) equilibrated in Buffer A. Peak fractions were combined, concentrated to <0.5 mL, and supplemented with 10% glycerol. Aliquots (10-50 μl) were flash-frozen and stored at −80°C. Typical stock concentrations ranged from 1–5 mg/mL.

### DNA binding assays

To generate dsDNA substrates ≤60 bp in length for binding assays, complementary synthesized ssDNA oligonucleotides (IDT Inc.) were resuspended in 10 mM Tris, 0.1 mM EDTA (pH 8.0). Equal amounts of each oligonucleotide (500 pmol; 5 μl of a 100 μM stock) were mixed in a final volume of 50 μl of water. The oligonucleotides were annealed in a thermal cycler using the following program: 95°C for 5 minutes, followed by a decrease of 0.5° C every 30 seconds for 170 cycles, holding at 25°C. Annealed dsDNA was purified using a DNA Clean and Concentrator kit (Zymo Research), eluting in 50 μl of 10 mM Tris, pH 8.5, quantified by Nanodrop and normalized to 500 nM. The dsDNA was assessed for purity on an 8% native acrylamide gel. Longer dsDNA substrates were generated by PCR amplification using Q5 Polymerase (New England Biolabs) and purified and analyzed as described above.

For DNA binding reactions, the DNA substrate was used at a final concentration of 50 nM. Untagged CasPRIB1 protein, diluted in Buffer A was added at final concentrations between 50 nM and 250 nM. 20 μl binding reactions were assembled in buffer containing 20 mM Tris, pH 7.9, 40 mM NaCl, 2.5% sucrose, and incubated at 22°C for 30 minutes.

Samples were resolved on 8% native acrylamide gels (29:1 acrylamide:bisacrylamide; Bio-Rad) containing 0.5 X TBE. Gels were pre-run for 15 minutes at 90 V in 1 X TBE, and wells were rinsed prior to sample loading. 5 μl of each binding reaction was loaded on the gel (no dye included) and run at room temperature for 70 minutes at 90 V in 1X TBE buffer. Following electrophoresis, gels were stained with SYBR Gold (Invitrogen) diluted in 1 X TBE for 30 minutes, rinsed twice in 1 X TBE, and imaged on a Bio-Rad system.

The percent binding for EMSA reactions was quantified by densitometry analysis of gels using ImageJ. Rectangles of equal size were used to quantify pixel density in the lower, unshifted band and the upper, shifted band for each reaction; adjacent rectangles were used to estimate background signal. After background subtraction, total DNA was calculated by summing the densities of both bands. The fraction of bound CasPRIB1 was then calculated as the ratio of shifted DNA to total DNA.

### Plaque assays

Lawns were generated by growing 5 mL cultures of *L. seeligeri* expressing type I-B CRISPR targeting ϕEGDe at 30°C overnight and combining 5 mL of molten BHI top agar with 5 μL of 5M CaCl2 and 100 μL of saturated overnight culture. Mixture was vortexed and poured over a BHI agar plate to solidify. 10-fold serial dilutions of phage were made in BHI and 2μL of each dilution were spotted on each top agar lawn. Plates were incubated at 30°C overnight for bacterial and phage growth.

---

**Figure S1.**
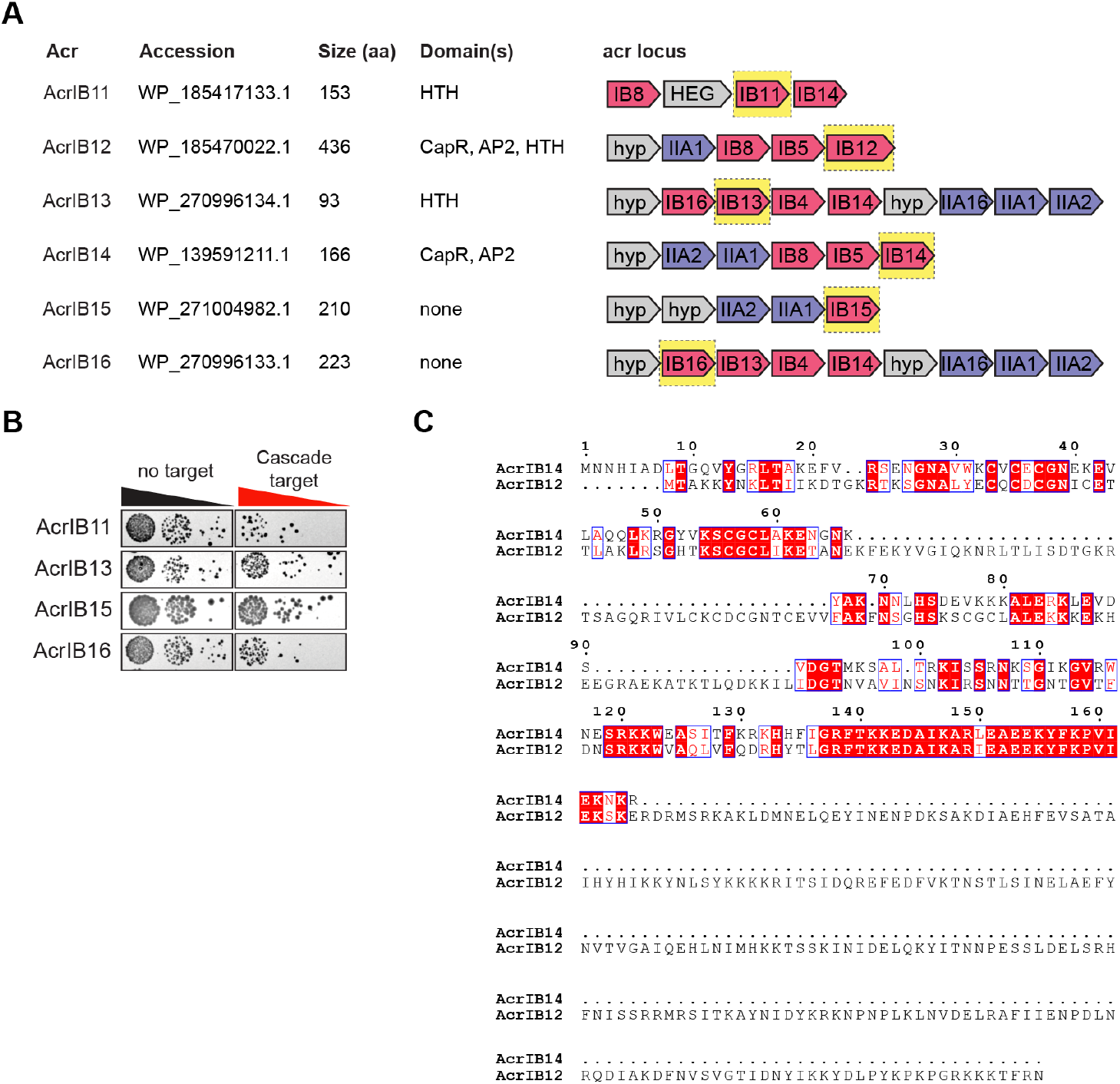
Novel type l-B Acrs. **(A)** Six novel Acrs from L. seeligeri phages and mobile genetic elements. Genetic context of tested acr gene sequences is depicted, with type l-B acr gene homologs in pink, type ll-Aacr homologs in blue. HTH, helix-turn-helix. **(B)** Plasmid targeting assay testing the activity of the indicated Acr on targeting of a conjugative plasmid by the L. seeligeri type l-B CRISPR-Cas system. Plasmid targeting assays for AcrlB12 and AcrlB14 are shown in Fig. 1C. **(C)** Alignment of AcrlB12 and AcrlB14 protein sequences.

**Figure S2.**
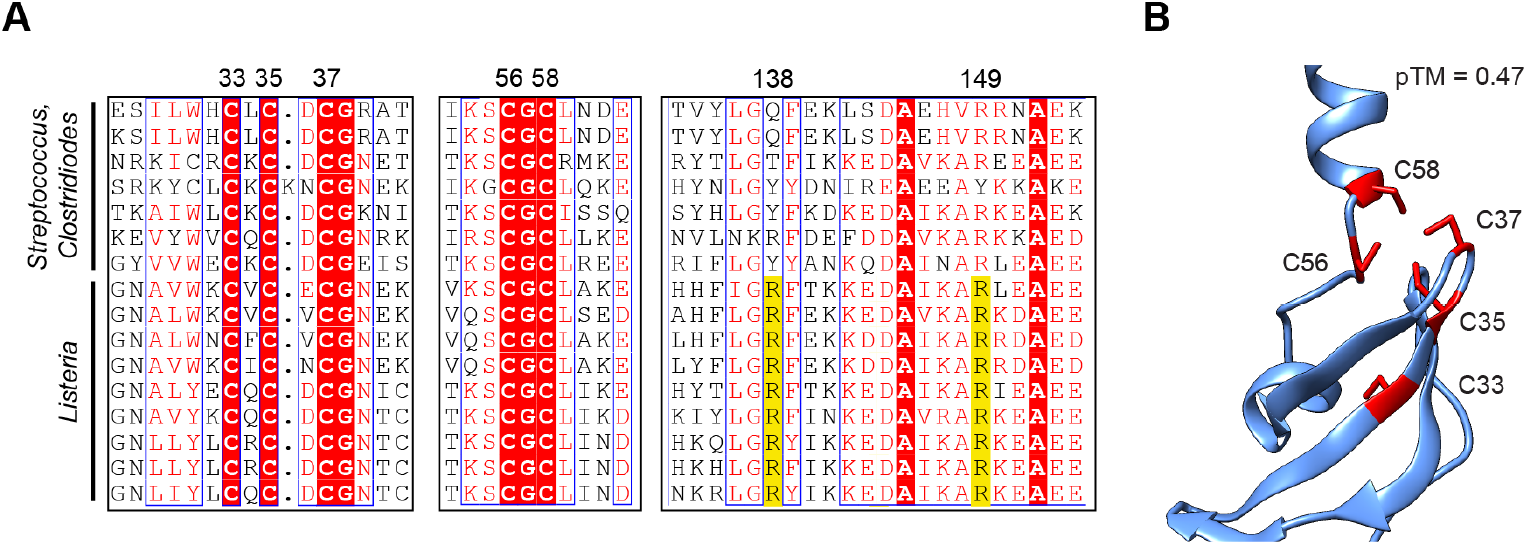
Conserved functional residues in the CapR and AP2 domains of CasPRs; **(A)** Multiple sequence alignment of 16 CasPR protein sequences showing conservation of cysteine residues in CapR domain, and arginine residues conserved in AP2 domain of Listeria representatives (yellow). **(B)** AlphaFold3 model of CasPRIBI, highlighting positions of conserved cysteines predicted to coordinate a Zn2+ ion.

**Figure S3.**
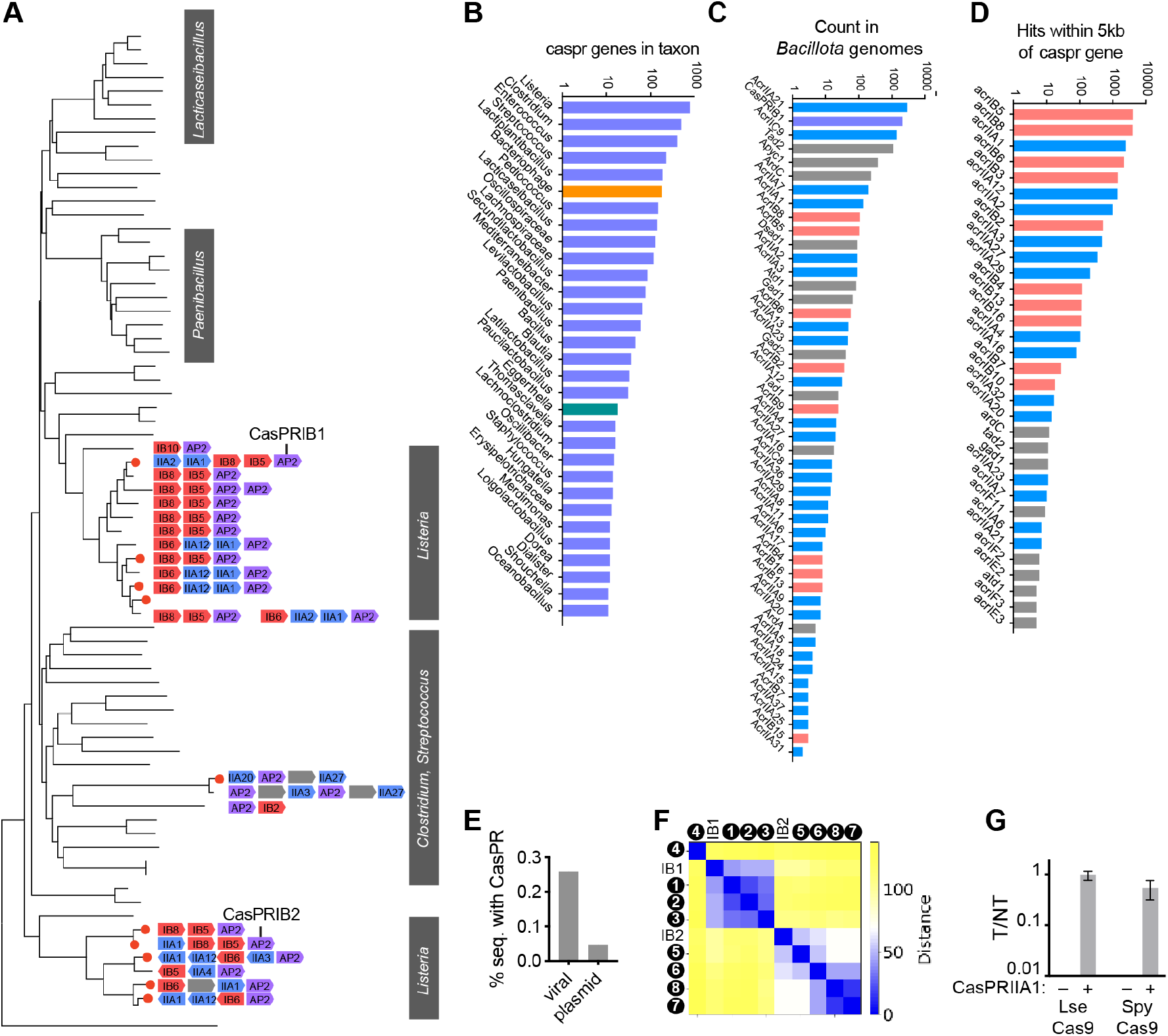
Bioinformatic analysis of CasPR homologs, related to Fig. 3. **(A)** Phylogenetic tree representing casPR homologs. Acr clusters encoding casPR homologs (violet) for each leaf are depicted. Homologs tested for activity are indicated with red cricles. Type l-B acr homologs in red, type ll-A in blue, gray genes indicate hypothetical proteins. **(B)** Occurrence of caspr genes in the indicated taxonomic groups. Taxa in blue are in phylum Bacillota, green in Bacteroidota, yellow are bacteriophages. L. seeligeri CasPRIBI was used as BLASTp query with <1E-4 E-value and >70% query coverage cutoffs. **(C)** Predicted defense inhibitor genes in Bacillota genomes. CasPRs (purple) and 137 other defense inhibitor proteins were used as queries in a TBLASTN search of the NCBI nt_prok database with <1E-4 E-value and >70% query coverage cutoffs. Type l-B Acr families (red), type ll-A Acr families (blue) and other defense inhibitors (gray) are indicated. **(D)** Defense inhibitor genes located nearby caspr genes. From the search described in (C), genes lying within 5000 bp of a caspr gene homolog were tabulated. **(E)** Percentage of sequences within viral (IMGVR) and plasmid (IMGPR) databases encoding CasPR homologs. **(F)** Distance matrix of tested CasPR homologs. **(G)** Plasmid targeting assay testing the activity of CasPRIIAI on target plasmid interference by Cas9 from *L seeligeri* (Lse) and Cas9 from *Streptococcus pyogenes* (Spy). T/NT, target to non-target transconjugant ratio. Error bars are SEM.

**Figure S4.**
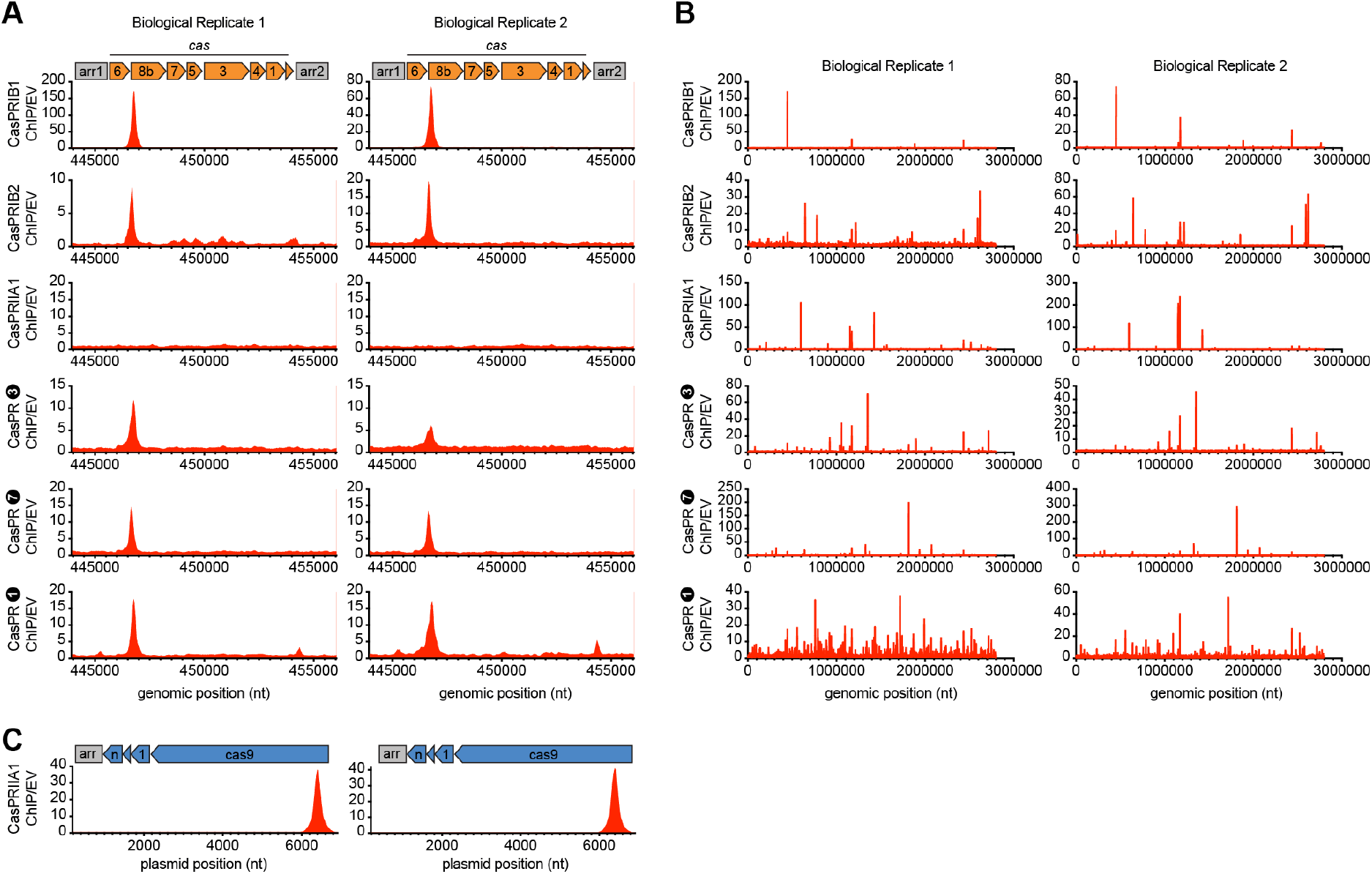
Site-specific binding of CasPRs to cas genes, related to Figure 4. **(A)** ChlP-seq profiles of indicated his6-tagged CasPRs, plotted along the LS1 type l-B CRISPR-Cas locus. Plot represents the ratio of reads in CasPR ChIP samples to those in empty vector (EV) control lacking CasPRs. ChIP signal was normalized to reads from a total input DNA sample. Two biological replicates are shown. **(B)** As in (A), but displaying CasPR ChIP profiles across the complete *L. seeligeri* LS1 genome. Two biological replicates are shown. **(C)** As in (A), but plotting the CasPRIIAI ChIP profile along the type ll-A CRISPR-Cas locus from L. seeligeri. Two biological replicates are shown.

**Figure S5.**
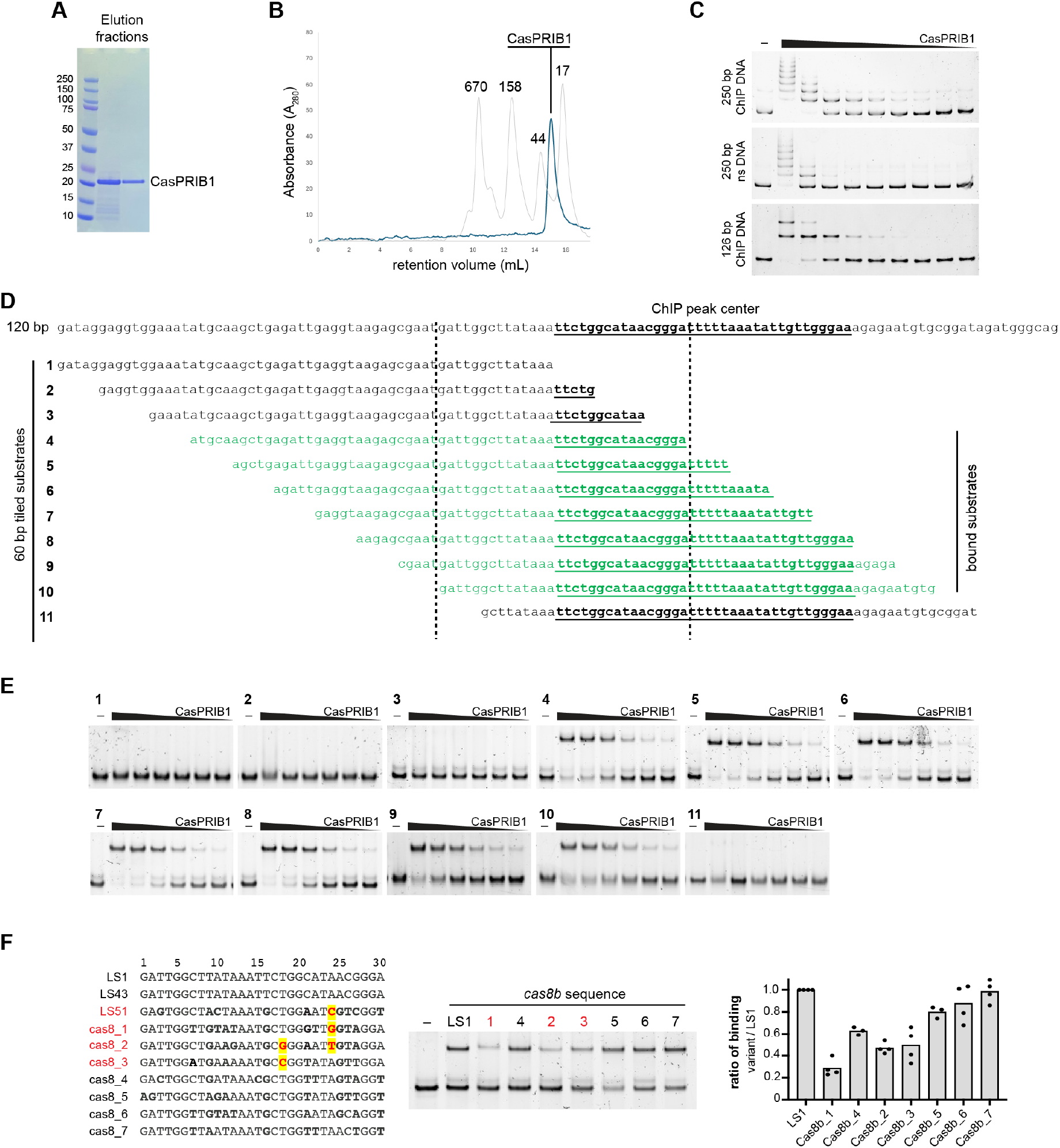
Identification of CasPRIBI recognition site, related to Figure 4. **(A)** Coomassie gel indicating purified *L. seeligeri* CasPRIBI sample, with molecular weight 19.1 kDa. Molecular weight marker in kDa. **(B)** Size exclusion chromatogram of CasPRIBI along with size standards. **(C)** EMSA with 50 nM of indicated DNA substate containing nonspecific (ns) or CasPRIBI ChIP sequence and 250, 125, 62, 31, 16, 8, 4, 2, 1, or 0 nM CasPRIBI. **(D)** WT 120 bp cas8b sequence bound by CasPRIBI, with ChIP peak center underlined, and twelve 60-bp tiled substrates, each overlapping by 5 bp, that collectively cover the interval. Substrates in green were bound by CasPRIBI in EMSA, indicating a region critical for binding (between dashed lines). **(E)** EMSA with 50 nM of indicated 60 bp DNA substrate (from panel D) and 0, 250, 200, 150,100, 75, 50 nM CasPRIBI, resolved on native acrylamide gels. **(F)** Seven diverse natural variants of the CasPRIBI binding site that vary from the site in LS1 at sites in bold were tested for binding in vitro. Substrates in red exhbited severely reduced binding. Quantification of variant binding relative to LS1 is shown for four replicates. Binding sites in strains LS43 and LS51, whose in vivo CasPR sensitivity is measured in Fig. 3E, are also shown.

## Notes

### Competing Interest Statement

AJM is a co-founder and advisor of Profluent Bio. The other author declares no competing interests.

